# The Core Promoter has an active role in interpreting patterning information

**DOI:** 10.64898/2026.10.11.758330

**Authors:** Susnata Karmakar, Kat Millen, Sarah Bray

## Abstract

Precise spatiotemporal control of gene expression is conventionally attributed to distal enhancers. The extent to which different core promoters actively shape transcriptional output from enhancers remains unclear, especially in the context of genes that contain alternate promoters. We have explored these questions using the *Drosophila grainy head* (*grh*) locus, and tested the activities conferred by its alternate promoters in combination with a well characterised Notch responsive enhancer. We find that, of the alternate promoters tested, only the *grh* short promoter is competent to drive neuroblast-specific expression. Notably, when paired with identical enhancers, *grh*-short and *grh*-medium promoters drive distinct expression patterns. The activity of other core promoters, such as hsp70, is also selective. Together the results establish that the core promoter acts alongside enhancers to confer specificity and robustness on gene expression profiles.

## Introduction

A central problem in developmental biology is how a limited repertoire of transcription factors and cis-regulatory elements generate the diversity of cell types that constitute complex tissues such as the nervous system. Cell-type specificity has largely been attributed to enhancers, distal cis-regulatory modules that integrate signalling and lineage inputs. Through the recruitment of transcription factors and cofactors these instruct when and where a gene is transcribed (Banerji et al., 1981; Palstra and Grosveld, 2012; Kawasaki and Fukaya, 2024). In contrast, the core promoter—the sequence surrounding the transcription start site that nucleates the preinitiation complex—has largely been treated as a passive endpoint that sets the level of transcription but not its specificity (Sikorski and Buratowski, 2009).

Yet core promoters are strikingly diverse. Firstly, distinct sequence elements, such as TATA box, Initiator element (Inr) and downstream promoter element (DPE) partition promoters into functional classes with different initiation properties and cofactor requirements (Haberle and Stark, 2018). These differ between developmental and housekeeping genes whose enhancers are intrinsically matched to the appropriate core-promoter class (Zabidi et al., 2015). In addition, promoter diversity can confer specificity on enhancer interactions, as certain enhancers activate one core-promoter class but not another even when located at equivalent distances (Butler and Kadonaga, 2001). Indeed transcriptional cofactors can themselves discriminate between core-promoter types (Haberle et al., 2019). Together these findings argue that core promoters can harbour regulatory information and participate actively in conferring specificity on gene transcription.

A further level of complexity arises with genes that carry multiple alternative promoters. Such arrangements are pervasive and deeply conserved. More than 40% of developmentally expressed *Drosophila* genes are transcribed from two or more promoters deployed in distinct, cell-type-specific patterns (Batut et al., 2013). The picture in mammals is similar, with alternative start sites accounting for most transcript-isoform differences across human tissues (Reyes and Huber, 2018) and many human promoters being composite, with multiple-start-sites that confer cell-type-specificity (Forrest et al., 2014). When, as in these cases, a single enhancer is presented with a choice of targets it is unclear how one promoter is selected over the others. Genomic proximity is one factor that can influence the choice, because enhancer-promoter (E–P) contact frequency has been found to decline with distance in some cases (Zuin et al., 2022). However, functional interactions routinely bridge large distances through chromatin looping (Yang and Hansen, 2024; Kittle and Levo, 2025) suggesting the absolute distance may not be a major factor. Furthermore, a single enhancer has been found to interact with four different gene promoters spread over a large genomic region (Masoura et al., 2025). The different patterns produced argue that the promoters help shape the identity and magnitude of the transcriptional response, rather than being passive players. Whether this is a widespread property requires further testing using systems where an enhancer is confronted with multiple native promoters.

*Drosophila grainy head* (*grh*) is an example of a locus that produces distinct tissue-specific isoforms. Grainyhead is the founding member of a conserved transcription-factor family that acts as a pioneer factor, engaging nucleosomal DNA to establish and maintain the accessible chromatin landscape of epithelial and other cell types (Jacobs et al., 2018; Nevil et al., 2020; Sundararajan et al., 2020). Its three vertebrate orthologues (GRHL1–3) are essential for epidermal barrier formation, craniofacial and neural-tube closure and inner-ear development. Their misregulation underlies human diseases ranging from orofacial clefting and neural-tube defects to hearing loss and epithelial cancers (Sundararajan et al., 2020). Across different species the use of alternative start sites appears to be a conserved mechanism to generate different functional isoforms of Grh-like proteins (Miles et al., 2017).

The eight transcripts that are produced from the *grh* locus fall into long, medium and short classes and initiate from several alternative promoters. In particular a neuroblast-specific protein isoform is produced through a combination of promoter and tissue-specific alternative splicing (Uv et al., 1997). It differs from the epidermal form in its DNA-binding preference and target repertoire, and its loss perturbs neural stem cell programmes and causes lethality (Almeida and Bray, 2005; Cenci and Gould, 2005; Uv et al., 1997). Neural expression of *grh* is regulated by Notch signalling, acting through defined, well-characterised enhancers (Almeida and Bray, 2005; Cenci and Gould, 2005; Wissel et al., 2018; Zacharioudaki et al., 2016). How the appropriate promoter is selected to produce the neural transcript—and hence which protein isoform is produced—is however not known. One question is whether there is a specific promoter compatibility driven by the signalling input from the Notch pathway.

Here we have investigated the functional characteristics of the different *grh* promoters to gain insights into the mechanisms governing enhancer-promoter compatibility. To achieve this, we paired individual native *grh* promoters first with the Notch responsive *grh* enhancer and subsequently with a range of other defined enhancers, testing their combined activity in transgenic flies. The distance from the enhancer was the same in all cases and, by using the same genomic landing site, all were exposed to the same genomic environment. Despite these conditions we found a clear difference in the activity of different promoters and most notably a single enhancer gave rise to distinct cell-type-specific patterns depending on the promoter with which it was paired. The *grh* short transcript promoter proved to be highly neuroblast-specific indicating it has an important role in conferring expression of the distinct isoform in these stem cells (Uv et al., 1997). Altogether the results demonstrate that core promoters are not simply permissive for transcription, they actively contribute to the tissue specificity and to the robustness of the transcriptional output, working together with the enhancer to confer distinct patterns of expression.

## Results

### Neuroblasts exclusively express *grh*-short isoforms

As a prelude to investigating mechanisms that enable tissue specific expression of *grh* isoforms, we first assessed which promoters are deployed in the larval CNS. The *grh* locus contains five alternative transcription start sites (ATSS) that produce eight distinct transcripts and protein variants (Fig. 1A). These can broadly be subdivided into three groups: *grh*-long, *grh*-medium and *grh*-short (Fig. 1A). Previous time-resolved analysis of larval central nervous system (CNS) transcriptomes (Wissel et al., 2018) suggested that the *grh* short isoforms are enriched in neuroblasts — the neural stem cells of the CNS (Fig. 1A).

**Figure 1.**
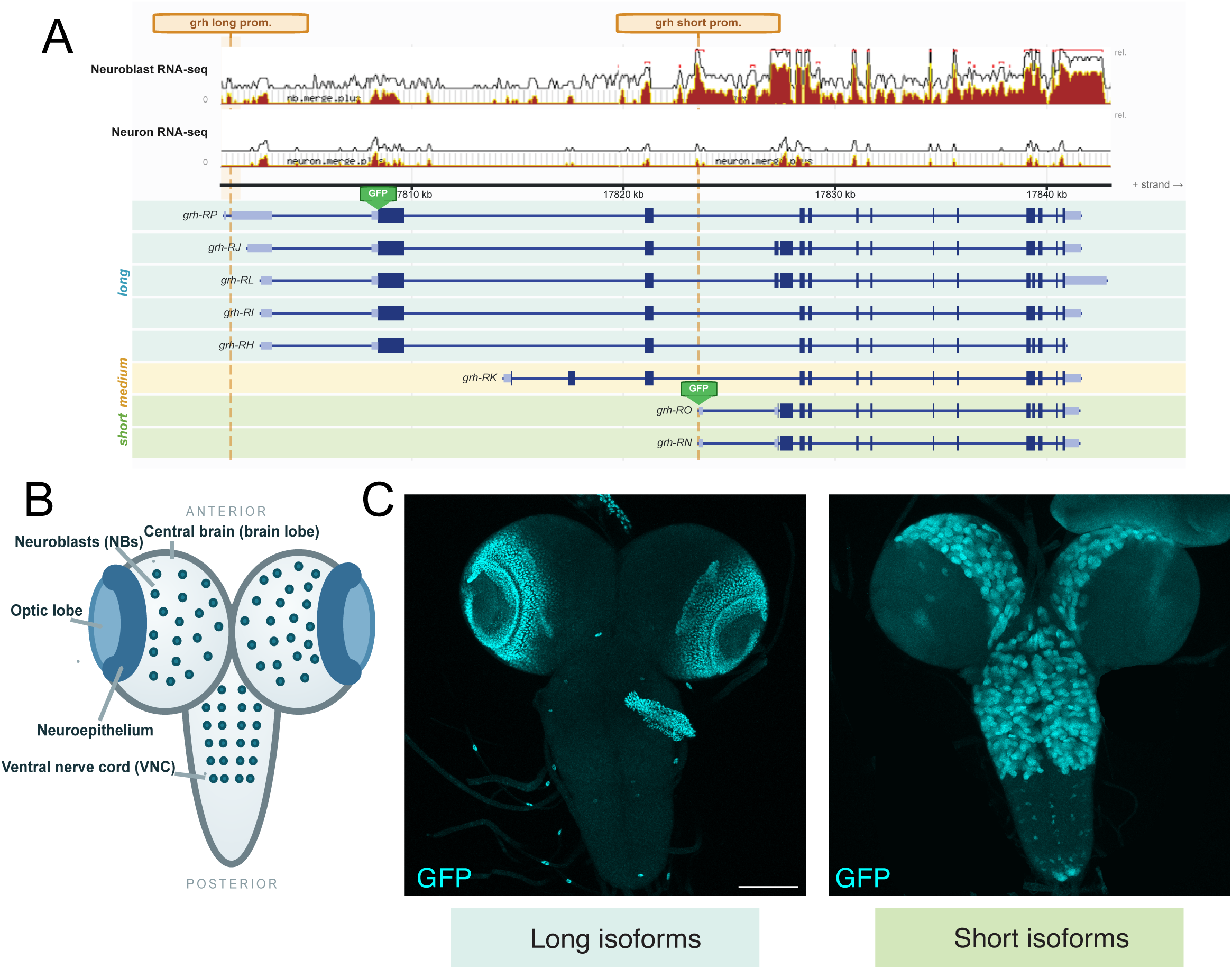
Long and short Grh isoforms have distinct expression in the larval CNS. (A) The *grh* locus (dm6, chr2R:17,801,000–17,843,000) with its eight annotated transcripts, grouped as long (*grh*-RP, -RJ, -RL, -RI, -RH; blue shading), medium (*grh*-RK, yellow shading) and short (*grh*-RO, -RN, green shading). Positions of long and short isoform promoters (brown) and sfGFP knock-in (green rectangle) are indicated. Upper graphs, neuroblast and neuron RNA-seq (Berger et al. 2012) (GSE38764). (B) Schematic of third-instar CNS with central brain, optic lobe, neuroblasts, neuroepithelium, ventral nerve cord indicated. (C) Expression (cyan) of long- and short-isoforms from GFP knock-in strains; maximum-intensity projections of third-instar central brain. Scale bars, 50 µm. Full genotypes are given in Table S3.

To characterise expression patterns of the “long” and “short” transcripts, we used engineered fly lines in which GFP had been inserted in frame into class-specific alternate exons. For *grh*-short, we used CRISPR-Cas9 to endogenously tag the *grh*-RN/RO first exon with GFP and for *grh*-long isoforms we utilized a previously generated fly line carrying sfGFP in frame within the first coding exon shared by all long isoforms (Nevil et al., 2020).

Strikingly, *grh*-long and *grh*-short had complementary patterns of expression (Fig. 1B, C; Fig. S1A). *grh*-short was robustly expressed in neuroblasts (NBs) and not in the neural epithelium (NE). In contrast, *grh* long isoforms were absent from the NB lineage, but robustly expressed in the NE (Fig. 1C). Similarly, short and long *grh* isoforms have distinct expression patterns in wing and leg discs, where the long isoforms are widely expressed through the epithelium whereas short isoform expression is restricted to a few locations, likely sensory lineages in these tissues (Fig. S1A).

If NBs exclusively express *grh* short-isoforms, as implied by the patterns observed, any mutations that specifically target the *grh*-short promoter should produce phenotypes similar to those observed for *grh*^370^, an allele that perturbs NB function (Almeida and Bray, 2005; Uv et al., 1997). Indeed flies harboring CRISPR generated mutations that disrupt *grh* short, *grh*^Δ15^ and *grh*^Δ10^ failed to execute the normal apoptosis of abdominal neuroblasts that occurs in late third-instar larvae (Fig. S1B), the phenotype previously described for loss of *grh* activity in this tissue (Almeida and Bray, 2005; Cenci and Gould, 2005). They also showed reduced viability when combined with null alleles for the *grh* locus (Fig. S1C).

Collectively, these results demonstrate that NBs exclusively express *grh*-short transcripts, indicating that the associated promoter is essential for expression in this cell-type, and hence that this isoform is responsible for the timely termination of some NB lineages (Fig. S1B, C).

### Promoters exhibit differential activity in combination with *grh* neuroblast enhancer

A fundamental question in transcriptional regulation is why one specific promoter is utilised when multiple promoters are available within a complex locus. To investigate the mechanisms governing promoter selection at the *grh* locus, we first combined each of the five native *grh* promoters with the previously defined Notch-responsive neuroblast enhancer (*grh[NB]*; (Zacharioudaki et al., 2016) to test their compatibility. In each case we selected a circa 150 bp fragment flanking the annotated transcription start-site (circa -100 to +50, see methods). Strikingly, the *grh*-short promoter was the only one of the five that was capable of driving robust reporter expression in NBs (Fig. 2B). In contrast reporters containing the other native promoters, which include one that is located closer to the enhancer in the genome (*grh*-medium), remained largely silent with little or no expression in NB lineages (Fig. 2B). In the absence of the enhancer, *grh*-short promoter alone resulted in no detectable reporter expression, confirming that it lacks autonomous enhancer-like activity (Fig. S2). Since the E-P distance was identical in all combinations tested, these results demonstrate that there is an intrinsic difference that governs promoter activity suggesting that their differential positions in the genome may not be a deciding factor in promoter selection.

**Figure 2.**
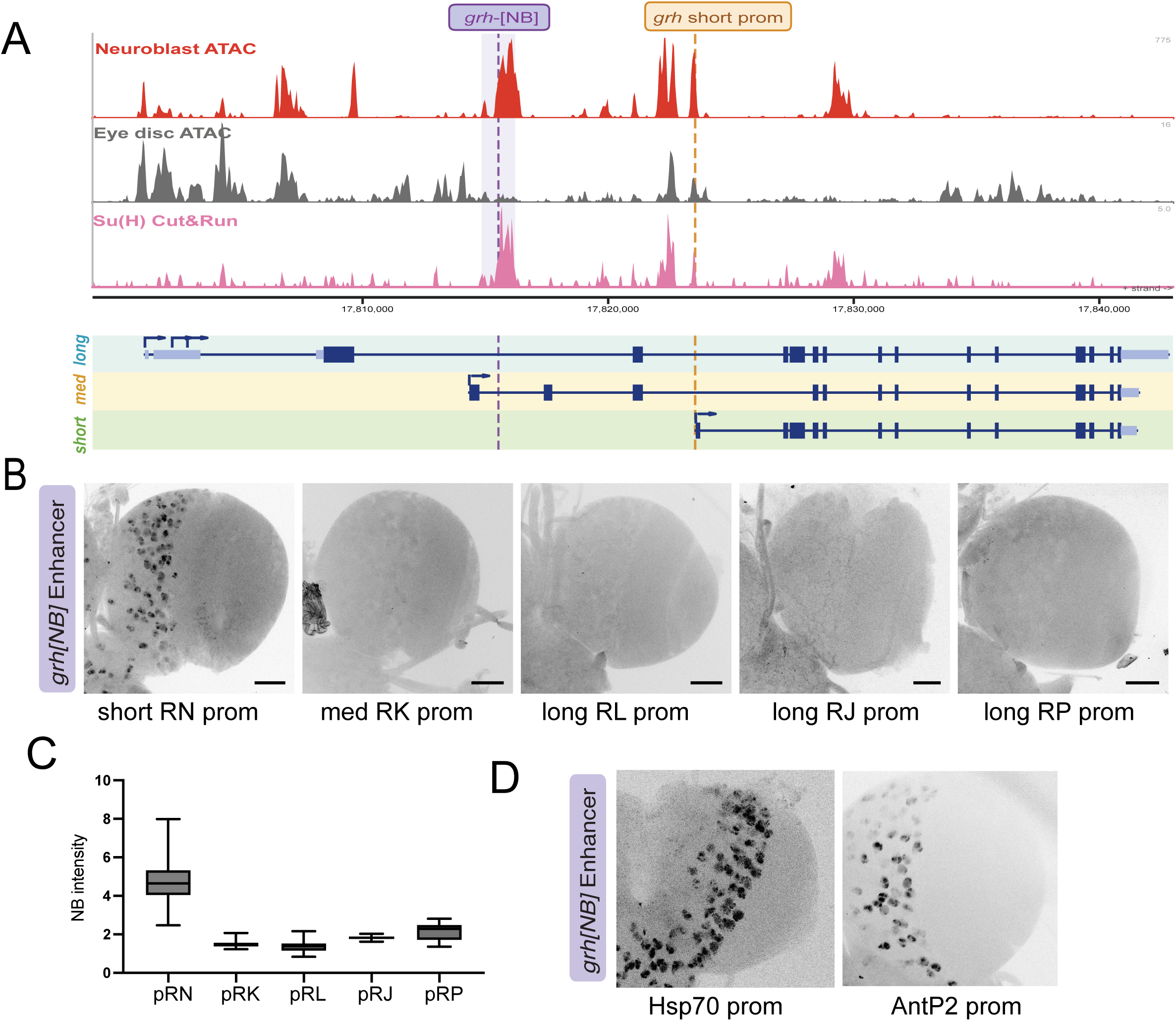
Only the short promoter fires with the *grh* neuroblast enhancer. (A) The *grh* locus with graphs depicting enrichments (log2) from neuroblast ATAC (GSM4561511; red), eye-disc ATAC (GSM8186965; grey) and Su(H) CUT&RUN (GSE218253; pink). *Grh*[NB] enhancer (purple; chr2R:17,814,846–17,816,211) and short promoter (orange; chr2R:17,823,564) are highlighted. (B) Expression in brain lobes driven by *grh*[NB] paired with each *grh* promoter (short RN, medium RK, long RL, RJ, RP); robust neuroblast expression is seen with *grh*-short (RN). Maximum-intensity projections, inverted greyscale; scale bars, 50 µm. (C) Quantification of *grh*[NB] reporter intensity from promoters as indicated, expressed as background-normalised GFP relative to local background (RN, n = 3 brains; RK, n = 2 brains; RL, n = 4 brains; RJ, n = 2 brains; RP, n = 4 brains). Data are plotted as boxes showing the mean ± s.d., with individual neuroblasts overlaid; groups were compared with an unpaired two-tailed Student’s t-test. (D) Expression in brain lobes driven by *grh*[NB] paired with *hsp70* (TATA) or *Antp-P2* (Inr) promoters; scale bars, 50 µm. Full genotypes are given in Table S3.

We next asked whether this activity arises from a unique requirement for specific sequences in the short promoter. To test this, we paired the *grh[NB]* with two non-native, widely used promoters: the TATA-containing minimal *hsp70* promoter (hsp70), also known as hs43, and the Initiator (Inr)-containing *Antennapedia* pAntp-P2 promoter. The first has been widely used in reporter constructs testing enhancer function and is generally assumed to be compatible with most enhancers (Hiromi and Gehring, 1987; Pfeiffer et al., 2008). In agreement, it conferred robust NB expression in combination with *grh[NB]* (Fig. 2D). The Inr-based pAntp-P2 promoter also yielded robust expression in neuroblasts when paired with *grh[NB]* (Fig. 2D). Thus, promoters with different core motifs can cooperate effectively with the *grh[NB]* enhancer. This behaviour is not restricted to the *grh*-short promoter.

Collectively, these findings demonstrate that of the 5 promoters in the locus, the *grh*-short promoter is uniquely capable of conferring NB expression in combination with the *grh[NB]* enhancer. As the promoters were all tested in the same configuration, this suggests that promoter sequences play an active role in determining expression outcomes, independent of physical distance.

### Promoter specificity occurs with a second Notch regulated enhancer

We next investigated whether the observed promoter selectivity was a unique property of interactions with the *grh[NB]* enhancer or was recapitulated with other enhancers. First, we utilised a Notch-responsive element (NRE) from the Deadpan (*dpn*) locus that contains three putative Suppressor of Hairless (Su(H)) binding motifs (Babaoğlan et al., 2013; San-Juán and Baonza, 2011) and paired it with the short (RN), medium (RK) and long (RL) *grh* promoters. In this context, the *grh*-short promoter remained the most efficient, enabling robust reporter expression in NBs. Unlike with *grh[NB]* enhancer, the expression domain included the NE, a region where the short *grh* isoform is not natively expressed (Fig. 3B). Of the other promoters, *grh*-med (RK), gave rise to moderate expression levels, most notably in the NE, while the long-isoform promoters produced no detectable expression (Fig. 3B, C).

**Figure 3.**
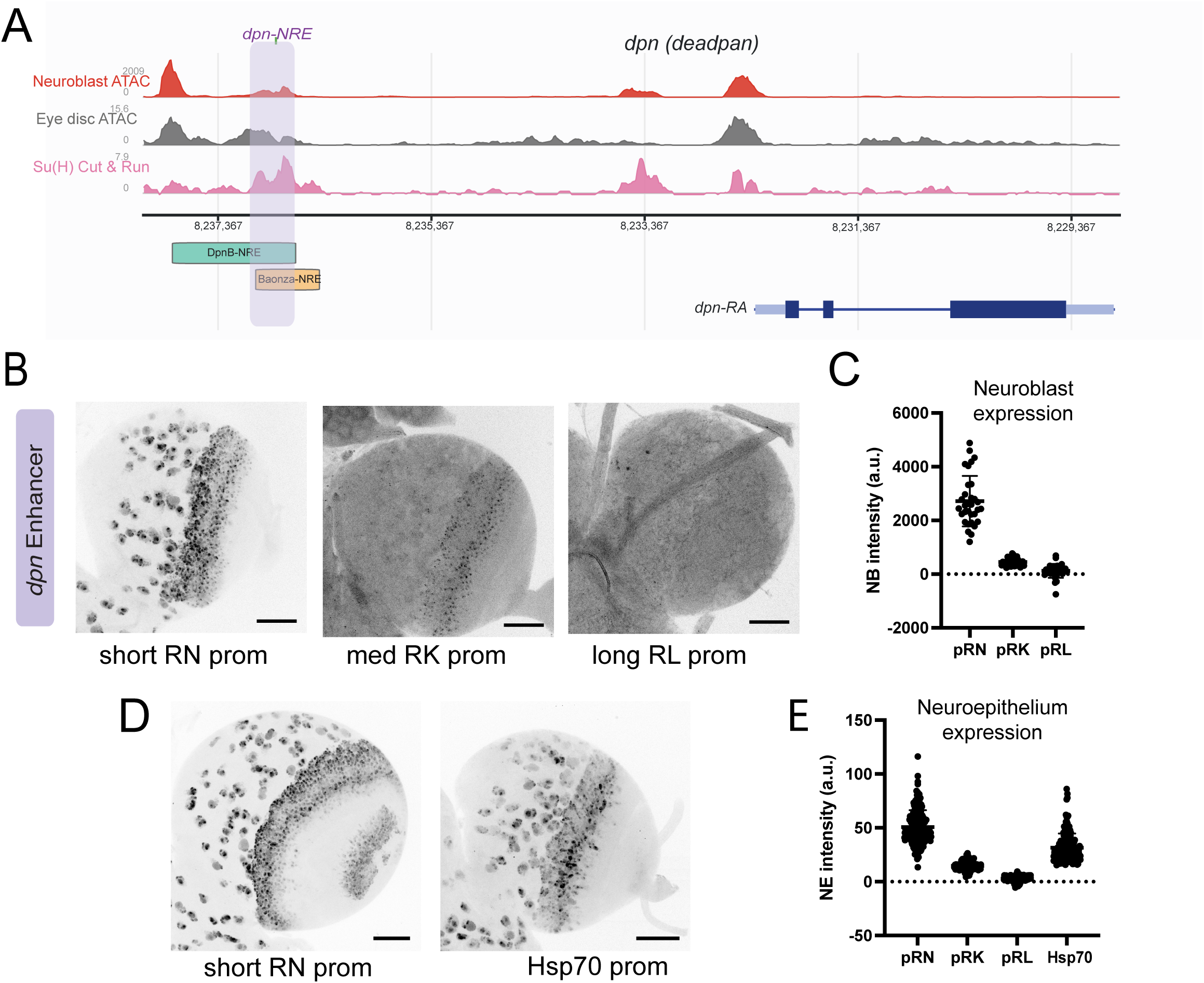
Promoter identity shapes the output of the *deadpan* neuroblast enhancer. **(A)** The *deadpan* (*dpn*) locus with graphs depicting enrichments in neuroblast ATAC (GSM4561511; red), eye-disc ATAC (GSM8186965; grey) and Su(H) CUT&RUN (GSE218253; pink); *dpn*[NRE] is highlighted (purple shading), and its relationship with previously defined *dpnB[NRE]* (teal) and *Baonza-NRE* (orange). (B) Expression in brain lobes driven by *dpn*[NRE] paired with *grh*-short (RN), *grh-*medium (RK) and *grh*-long (RL) promoters. Maximum-intensity projections, inverted greyscale; images within a panel were acquired in a single session at matched gain; scale bars, 50 µm. (C) Quantification of neuroblast expression intensity driven by *dpn*[NRE] with promoters as indicated (RN, n = 42 neuroblasts from 3 brains; RK, n = 32 from 3; RL, n = 31 from 2). Data are plotted as boxes showing the mean ± s.d., with individual neuroblasts overlaid (D) Expression in brain lobes driven by *dpn*[NRE] with *grh*-short (RN) or *hsp70* promoters; scale bars, 50 µm. (E) Quantification of neuroepithelium expression intensity driven by *dpn*[NRE] with promoters as indicated (RN, n = 3 brains; RK, n = 3 brains; RL, n = 2 brains; hsp70, n = 3 brains). Data are plotted as boxes showing the mean ± s.d., with individual neuroblasts overlaid; groups were compared with an unpaired two-tailed Student’s t-test. Full genotypes are given in Table S3.

We compared the activity produced by *dpn[NRE]* when paired with the basal *hsp70*. In this case the profile was similar to that of *grh*-short, with high expression in both NBs and NE (Fig. 3D, E). These results demonstrate that the outcome from the enhancer is modified depending on the promoter with which it is paired. In particular, *grh*-med promoter can only support robust expression in the NE and not in the NBs. We note also that *grh*-short promoter behaves in a very similar manner to *hsp70* despite the fact that it lacks any obvious TATA-element.

We further evaluated the performance of *grh*-short in the developing photoreceptors of the eye imaginal disc, where *dpn[NRE]* is also regulated by Notch signaling (Mavromatakis and Tomlinson, 2016). Both *grh*-short and *hsp70* promoters yielded similar expression patterns in the emerging photoreceptors, with *grh*-short producing a more even and robust pattern than *hsp70* (Fig. S3). This confirms that *grh*-short is able to function robustly in tissues where it is not usually utilised, since the Grh-short protein isoform is not expressed there (Fig. S1A).

Taken together, these observations demonstrate that the promoter is more than a passive entity and plays an instructive role in the patterns of expression as well as in the levels of transcription produced (Fig. 3, Fig. S3).

### *grh* promoters confer cell-type-specificity on more general enhancers

Both enhancers tested are Notch signalling responsive. We therefore wondered whether the promoter compatibility was linked to the fact that these are signalling-responsive and tested this using more ubiquitously active enhancers. We therefore selected the *β-Tubulin 56D* enhancer (*βTub56D)*, which is broadly active across the larval CNS (Buttgereit, 1993; Buttgereit et al., 1991), and the widely expressed *Actin 5C* enhancer (*Act5C*; Minervini et al., 2010).

The *grh*-short promoter was highly efficient in combination with *β-Tubulin 56D* enhancer, producing robust reporter expression in NBs, some early progeny and the NE (Fig. 4B; Fig. S4). Most strikingly, *grh*-med promoter generated a unique pattern, with moderate expression in differentiated neurons, as well as in the neuroepithelium. However, no expression was detected in the NBs. The *hsp70* promoter conferred a mixture of the two responses, with robust expression in NBs, NE and some neurons. Thus, a single enhancer generates distinct expression patterns depending on which of the promoters it is paired with (Fig. 4B). Core promoters can thus govern cell-type-specific transcriptional outcomes.

**Figure 4.**
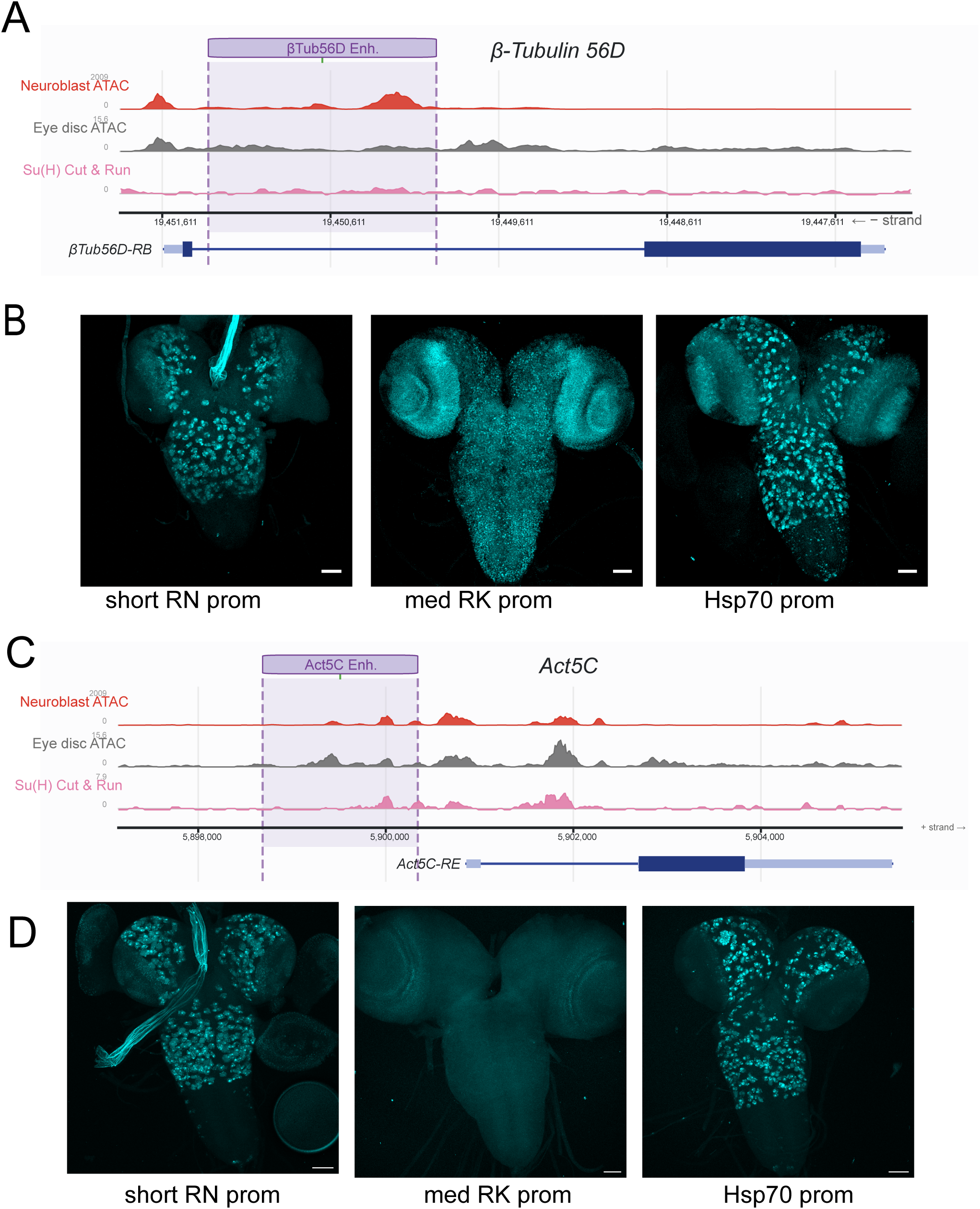
Promoter identity restricts the output of signalling-independent enhancers. (A) *β-Tubulin 56D* locus with enrichment tracks as in Figure 3, the *βTub56D* enhancer is highlighted (purple). (B) Whole CNS expression (cyan) driven by *βTub56D* enhancer paired with *grh-*short (RN), *grh*-medium (RK) and *grh*-long (RL) promoters (scale bars, 50 µm; gain on panels is not equivalent). (C) *Actin 5C* locus (chrX) with the enrichment tracks as in Figure 3 and *Act5C* enhancer highlighted (purple). (D) Expression from *Act5C* enhancer paired with *grh*-short (RN), *grh*-medium (RK) and *hsp70* promoters; scale bars, 50 µm. Full genotypes are given in Table S3.

The *grh*-short promoter exhibited the same robust expression pattern in NBs and their immediate progeny and in the NE when combined with the *Act5C* enhancer (Fig. 4D; Fig. S4). In contrast, the *grh*-med promoter remained inactive in NBs even with this ubiquitous enhancer, but it gave rise to a distinct pattern in other parts of the CNS. Notably, this was significantly different from the pattern observed with the *βTub56D* enhancer arguing for a complex interplay between the enhancer and promoter sequences that can generate unique cell specificities (Fig. 4D).

Collectively, these findings demonstrate that core promoter sequences play an active role in the expression profiles that are produced by enhancers.

### Features of the *grh*-short promoter that contribute to activity

Having found that *grh*-short is uniquely competent among the *grh* promoters, we asked what sets it apart. First, we searched all five *grh* promoters for matches to known core-promoter motifs (Materials and Methods). Neither *grh*-short nor any of the other *grh* promoters contain a match to the well characterised TATA-box motif. However, *grh*-short contains an Inr element close to the transcription start site and a region with homology to the DPE (Fig. 5A). These motifs are recognised by transcription factor II D (TFIID) through Taf6 and other subunits (Burke and Kadonaga, 1997; Louder et al., 2016). In agreement depletion of *Taf6* in NBs (with the neuroblast-restricted *grh*-GAL4) resulted in reduced levels of expression (Fig. 5B,C). However, although absent from the three *grh*-long promoters the Inr/DPE core configuration is also present in *grh*-med and so cannot explain the distinct function of *grh*-short.

**Figure 5.**
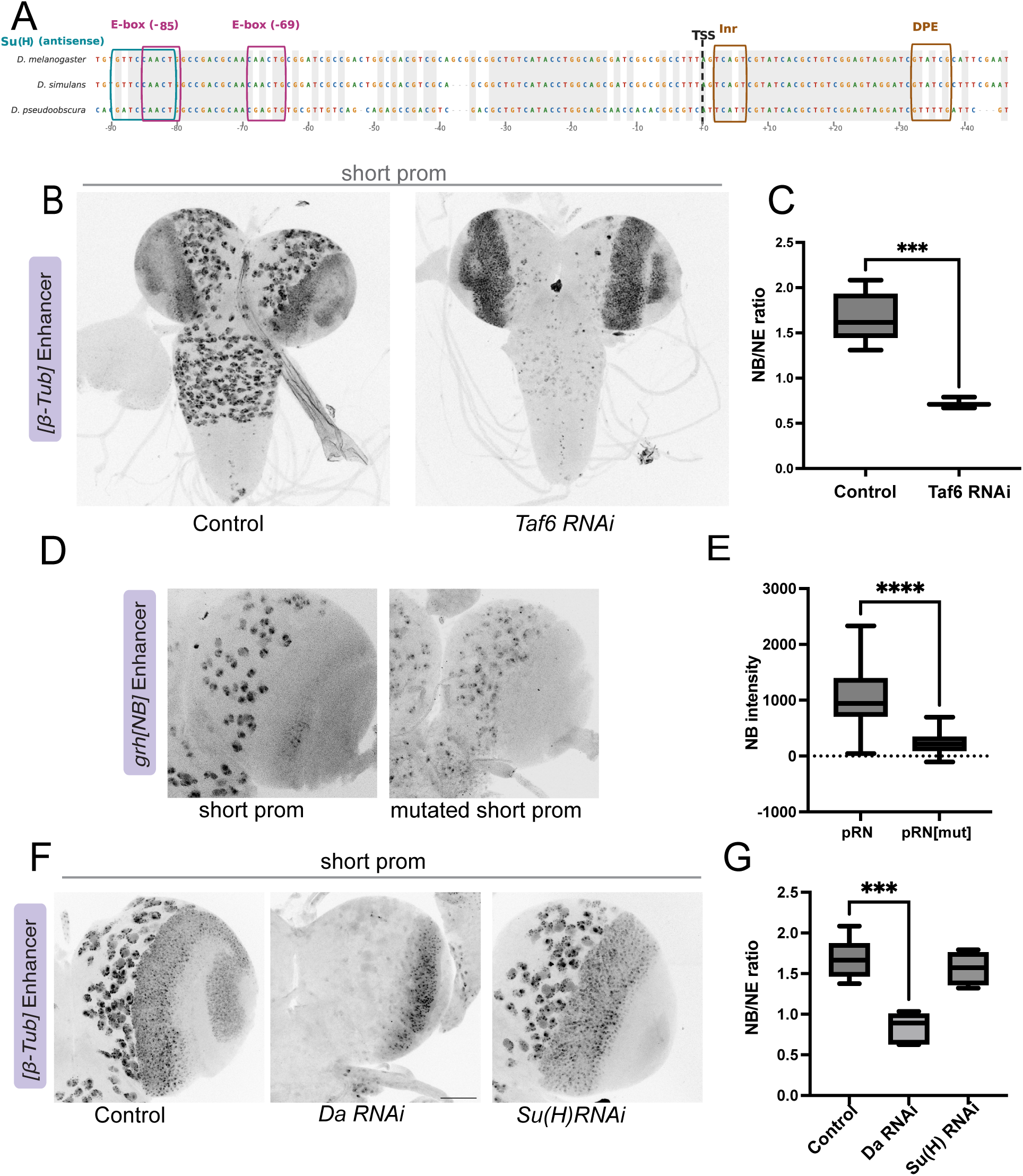
The short promoter depends on TFIID and on a conserved proximal proneural-bHLH element. (A) Cross-species alignment of *grh*-short (pRN) proximal region (≈−98 to +50 bp; *D. melanogaster*, *D. simulans*, *D. pseudoobscura*), marking the conserved proximal motif (the 12 bp scrambled in the reporter), with an overlapping Su(H)/CSL motif (cyan), and E-boxes (−85, and −69; magenta). Initiator (Inr; TCAGT) and DPE motifs also marked. (B) Expression in CNS from *βTub56D* enhancer in combination with *grh*-short in control versus *Taf6* RNAi; Taf6 knockdown reduces neuroblast expression. Scale bars, 50 µm. (C) Quantification of neuroblast-to-neuroepithelium expression levels, *lacZ* control versus *Taf6* RNAi. Because *grh*-GAL4 drives the RNAi in neuroblasts only, while the reporter is also expressed in the neuroepithelium, the neuroepithelial signal serves as an internal normaliser (*UAS-lacZ* control, n = 6 brains; *Taf6* RNAi, n = 3 brains). Data are plotted as boxes showing the mean ± s.d., with individual brains overlaid, the NB/NE ratio being a per-brain measurement; groups were compared with an unpaired two-tailed Student’s t-test (P value 0.0002). (D) Expression in brain lobes from *grh[NB]* paired with wild-type (RN) and mutated (RNmut) *grh*-short. Scale bars, 50 µm. (E) Quantification of neuroblast reporter intensity (RN, n = 4 brains; RN[mut], n = 4 brains). Data are plotted as boxes showing the mean ± s.d., with individual neuroblasts overlaid; groups were compared with an unpaired two-tailed Student’s t-test (P < 0.001). (F) Expression in brain lobes driven by *βTub56D* enhancer in combination with *grh*-short in control, *daughterless* (*da*) RNAi and *Su(H)* RNAi conditions. Scale bars, 50 µm. (G) Quantification of the neuroblast/neuroepithelium ratio: control versus *da* RNAi (P < 0.001) and *Su(H)* RNAi (n.s.) (UAS-lacZ control, n = 6 brains; *da* RNAi and *Su(H)* RNAi, n = 4 brains each). Data are plotted as boxes showing the mean ± s.d., with individual brains overlaid, the NB/NE ratio being a per-brain measurement; groups were compared with an unpaired two-tailed Student’s t-test (P < 0.001). Full genotypes are given in Table S3.

The *grh*-short promoter region is moderately conserved across *Drosophila* (Fig. 5A). Searching for transcription factor (TF) binding motifs within regions of conservation we identified an imperfect match to a Su(H) binding motif. As there is detectable Su(H) binding at *grh*-short in NBs (Fig. 2A) this raised the possibility that the promoter motif could mediate E-P interactions between Notch regulated *grh[NB]* enhancer and *grh*-short. To investigate, the 12 bp containing the motif were scrambled and the activity of the resulting mutated promoter (*grh*-short[mut]) assessed. The expression levels within the NBs were markedly reduced and there was no change in specificity suggesting the mutated region contributes to *grh*-short activity (Fig. 5D, E). However, a similar reduction in levels occurred when *grh*-short[mut] was combined with the *βTub56D* and *Act5C* enhancers (Fig. S5A–D), arguing that the motif’s role is not specific to Notch-regulated enhancers but is more generally linked to the promoter efficiency. Furthermore, introducing the same 12 bp into the *grh*-long (RL) promoter did not confer NB activity (Fig. S5E, F).

Two approaches were taken to identify other TFs that might act on *grh*-short. First, the 12 bp scrambled in *grh*-short[mut] were surveyed for matches to known TF binding motifs. This identified an E-box motif associated with Daughterless (Da)/proneural bHLH (CAGCTG; JASPAR MA2308.1) which overlaps the Su(H) motif (Fig. S6D). Second, we asked which TF motifs were enriched in *grh*-short and *dpn* promoters, but not in *grh*-med or *grh*-long (RL). Candidates from both approaches were then cross-referenced against expression data to identify those expressed in NBs, yielding a short-list of eight candidates. To evaluate the effects of depleting these TFs on expression from *grh*-short promoter, we tested it in combination with the *βTub56D* enhancer (Fig. S6A). As this combination gives rise to expression in NBs and NE, and the knock-down was restricted to the NBs, the expectation was that any expression changes should be present in the NBs only. Of the factors tested, only Da fulfilled this criterion: depletion of *da* reduced expression from *grh*-short within the NBs while leaving the NE signal unchanged, whereas depletion of Su(H) had no significant effect (Fig. 5F, G). The *Su(H)* RNAi line was itself effective, since it strongly reduced a Su(H)–GFP reporter in the wing disc (Fig. S6B, C), suggesting that the imperfect Su(H) motif in *grh*-short is unlikely to be a functional CSL site. Thus, Da emerges as a bona fide candidate to confer activity on *grh*-short, since both the knock down of *da* and the scrambling of its binding motif within the *grh*-short promoter were sufficient to compromise expression.

## Discussion

Testing the five native promoters of a single locus against a common set of enhancers, we find that a given enhancer can yield different cell-type-specific outputs depending on which promoter it engages. Of those tested, only *grh*-short is competent in neuroblasts. Promoter identity therefore informs where a gene is expressed, which isoform is transcribed and what levels of RNA are produced. In the case of *grh* this imposes an additional layer of control upstream of the tissue-specific splicing previously described at this locus (Uv et al., 1997).

### Compatibility can override proximity

Since all promoters were tested against the same enhancer, at a common landing site and at the same distance from it, the differences between them cannot be attributed to their relative positions or to other restrictions imposed by genomic conformations, such as TADs. Perhaps one of the most striking results is the different cell-type profiles conferred by the heterologous *βTub56D* enhancer with *grh*-short and *grh*-med promoters. The resulting cell-specific profiles illustrate that the output from a given enhancer is profoundly dependent on the promoter with which it is paired. In this context, the promoter appears to gate the activity of the enhancer to channel it in one or other cell type.

A second notable finding is that the three *grh*-long promoters were inefficient with every enhancer tested and in all tissues examined. They are also lacking sequence matches with any of the recognised core-promoter classes (Ohler et al., 2002; Juven-Gershon and Kadonaga, 2010). At face value, this might suggest that they are at best weak promoters. However, the expression levels of the knock-in line that captures expression of the long isoforms argues against this simple interpretation, as there is robust expression detected at all developmental stages (e.g. Fig. 1C, Fig. S1A). Their inactivity in the reporter assays may therefore be a property of their inability to operate in combination with the enhancers tested. This would imply a requirement for specific enhancer properties that are present in their native enhancers but not in those tested. This type of specificity may be important in the selection of one promoter over another in vivo (Balasubramanian et al., 2024).

The concept of enhancer–promoter compatibility is also relevant for the interaction between the *grh[NB]* enhancer and *grh*-short promoter. In the gene locus, *grh[NB]* is in close proximity to *grh*-med but >7 kb from *grh*-short, (Fig. 1), yet it operates exclusively with the latter to produce the *grh*-short transcripts in neuroblasts. This specificity is recapitulated in the reporter combinations demonstrating there is a fundamental difference in promoter compatibility that likely underpins the usage in the native context. This appears to be a more general property as such specificity is seen with other promoter combinations (Masoura et al., 2025).

### Features of promoter compatibility

By supporting the assembly of the pre-initiation complex, core promoters guide transcription initiation from precise positions at defined levels (Juven-Gershon and Kadonaga, 2010). Their sequence composition plays a central role in setting gene expression levels, based on mutational analyses of synthetic promoters (Qi et al., 2022). However, evidence argues against the differences in *grh* promoter specificity being solely attributable to effects on levels. For example, while *grh*-short confers robust expression in the neuroblasts and in the eye disc in combination with *dpn[NRE]*, when paired with *Act5C* enhancer it only supports expression in the CNS and not in the eye disc. This argues for a more complex regulatory relationship between the enhancers and promoter.

A further feature of promoters that confers a degree of selectivity is the type of core motifs that they contain. At least four different core promoter structures have been identified which affect levels of expression and make preferential interactions with different types of regulatory elements (Butler and Kadonaga, 2001; Zabidi et al., 2015; Haberle et al., 2019; Qi et al., 2022). However, our *in vivo* observations indicate that core-promoter structure cannot fully account for the different capabilities. Both *grh*-short and *grh*-med have an Inr/ DPE but only the former yields robust neuroblast expression with *grh[NB]*. Similarly, *grh[NB]* can work as efficiently with the TATA-containing hsp70 as with the Inr-containing *grh*-short and *Antp*-P2 promoters (Fig. 2D). The discrimination among the promoters thus extends beyond the core features captured by the classifications. Evidence for discriminatory behaviour has also emerged from massively parallel assays measuring enhancer–promoter combinations (Martinez-Ara et al., 2022).

As the main task of core promoters is to support the assembly of the pre-initiation complex, many of the conserved features are associated with recruitment of TFIID, as for the INR/DPE in *grh*-short. Large-scale analyses have in addition identified some commonly occurring motifs, which include the E-box, a DNA motif that binds bHLH transcription factors and was found to peak in both *Drosophila* and human promoters (FitzGerald et al., 2006; Qi et al., 2022). An E-box, along with a partial match to a Su(H) motif, were among the potential regulatory inputs in a conserved region of *grh*-short. Scrambling this lowered the output, which was recapitulated by the depletion of the bHLH factor daughterless but not by depletion of Su(H). Thus it appears that E-box binding factors make an important contribution to promoter activity levels, but in our experiments there was no evidence that they confer specificity on enhancer promoter interactions or cell-type activity.

### Evolutionary advantages of Promoter specificity

Although questions remain about what makes a promoter competent to respond to a given enhancer, our results argue that promoters can play a substantial role in determining which transcripts are produced at a given time and location. In this manner, the acquisition of alternative promoters introduces greater regulatory and functional diversity without gene duplication. When loci accumulate alternative promoters and enhancers, as occurred with *grh* (Kuzin et al., 2018; Sipani et al., 2025), their compatibility will be a key factor in determining how the gene responds to different inputs and in turn what the functional consequences will be.

## Materials and Methods

### Drosophila stocks and husbandry

Flies were reared on standard cornmeal medium and all crosses were performed at 25 °C unless otherwise stated. The stocks used are listed in Table S1. *grh*^Δ10^ and *grh*^Δ15^ short-promoter deletion alleles were generated in this study (see *CRISPR/Cas9 genome editing*); *grh*^B37^ and *Df(2R)Pcl7B* were described previously (Duncan, 1982; Bray and Kafatos, 1991).

### CRISPR/Cas9 genome editing

Endogenous sfGFP knock-in. A superfolder GFP (sfGFP) coding sequence (717 bp) was inserted in frame at the first exon shared by the *grh* short (RN/RO) isoforms by homology-directed repair. Two guide RNAs flanking the first exon were used (gRNA-A, GACTGGCGACGTCGCAGCGG; gRNA-B, GGCAGTAGTGCAATATTCTG). A scarless donor was assembled by NEBuilder (Gibson) assembly from a pHD-scarless-DsRed backbone (Gratz et al. 2014) together with the left homology arm, an exon1–sfGFP fragment, a 3×P3–DsRed screening cassette and the right homology arm; PAM sites within the donor were mutated to prevent re-cleavage and the translation-initiation context was optimised. Correct integration at the *grh* locus was validated by PCR with primers anchored outside the homology arms and by sequencing. The long-isoform sfGFP line was as described previously (Nevil et al. 2020).

*grh* short-promoter deletion alleles. To disrupt the short promoter and its first exon, the same two guide RNAs (together spanning a 311 bp region encompassing the RN first exon) were cloned into pCFD4 (Port et al. 2014) and injected into a *nos*-Cas9 strain. Two independent deletion alleles, *grh*^Δ10^ and *grh*^Δ15^, were recovered; in each the RN first exon is deleted between the two cut sites and a short non-templated sequence is retained at the repair junction (41 bp in *grh*^Δ10^ and 15 bp in *grh*^Δ15^), disrupting the short-isoform reading frame. Lesions were confirmed by Sanger sequencing.

### Reporter constructs

Enhancer–promoter reporter constructs were built in a modified pBDP backbone driving an mNeonGreen reporter (Ceolin et al., 2020). Unique *Hind*III, *Spe*I, *Bgl*II and *Eco*RI sites were introduced by assembling a synthesised insert into the *Bgl*II/*Eco*RI-linearised vector, so that enhancers could subsequently be cloned between the *Hind*III and *Spe*I sites and promoters between the *Spe*I and *Eco*RI sites; fragments were joined by restriction digestion and ligation or by Gibson assembly. For each *grh* isoform, the promoter was defined as 100 bp upstream to 50 bp downstream of the annotated transcription start site (≈150 bp fragment; see Table S2). Promoter fragments (short RN; medium RK; long RL, RJ, RP) were introduced as synthesised double-stranded DNA or as annealed oligonucleotides carrying *Spe*I/*Eco*RI-compatible overhangs; the non-native *hsp70* (hs43) and *Antp-P2* promoters were included for comparison. The *Antp-P2* fragment was amplified from genomic DNA to the same −100/+50 design (dm6 chr3R:6,932,791–6,932,940, minus strand), and the hs43 cassette, which comprises the 43 bp immediately upstream of the *hsp70* transcription start site together with the first 84 bp of its 5′ leader, was amplified from an existing reporter plasmid (Table S2). Details of enhancer fragments *grh*[NB]-NRE, *dpn*[NB]NRE (amplified with flanking *Hind*III/*Spe*I sites), *β-Tubulin 56D* (the AS1 region, ≈1.36 kb, Buttgereit, 1993) and *Actin 5C* (Act5C-RE, FBtr0345894, Minervini et al. (2010)) are provided in Table S2. All constructs were verified by Sanger sequencing.

### Site-directed mutagenesis

To test the predicted proximal E-box/Su(H) element in *grh*-short promoter, the 12 bp proximal motif TTCCAACTGGCC (dm6 chr2R:17,823,477–17,823,488; −88 to −77 relative to the *grh*-short transcription start site at chr2R:17,823,565) was replaced by ATACTCATAGAA, the sequence occupying the equivalent position in the long (RL) promoter, which abolishes the E-box match; in the reciprocal pRL-mod construct TTCCAACTGGCC was introduced at the equivalent position of the RL promoter, −88 to −77 of the RL transcription start site. In each case the native flanking sequence of the promoter was left unchanged. The replaced element contains the conserved −85 E-box (CAACTG; chr2R:17,823,480–17,823,485) and overlaps the imperfect Su(H)/CSL match; a second E-box at −69 (CAACTG; chr2R:17,823,496–17,823,501) lies outside it and was left intact. Both modified promoters were synthesised as 150 bp double-stranded fragments carrying SpeI- and EcoRI-compatible overhangs and cloned into the SpeI/EcoRI-digested *grh[NB]* backbone by Gibson assembly, generating *grh[NB]*-pRN[mut] and *grh[NB]*-pRL[mut]; the modified short promoter was also combined with the βTub56D and Act5C enhancers. Promoter fragments and their coordinates are listed in Table S2, and all constructs were confirmed by sequencing.

### Transgenesis

Reporter constructs were integrated at the *attP40* landing site (chromosome 2L, cytological position 25C) by φC31 integrase-mediated recombination. Plasmids were injected into embryos of Bloomington *Drosophila* Stock Center #25709 (*y^1^ v^1^ P{nanos-phiC31\int.NLS}X; P{CaryP}Msp300^attP40^*), which provides *nanos*-driven φC31 integrase and the *attP40* site; Surviving G_0_ adults were crossed to *y w*, and transgenic progeny were identified by red eye colour (mini-white marker). Independent transformants were then balanced over second-chromosome balancers to establish stable stocks.

### RNAi

UAS-RNAi expression was driven with *grh*-GAL4, which is active in neuroblasts; a UAS-*lacZ* transgene served as the control, and crosses were maintained at 25 °C. Taf6 was depleted using BDSC #57181. The candidate screen tested *klumpfuss* (*klu*, BDSC_28731), *broad* (*br*, BDSC_51189), *worniu* (*wor*, BDSC_77364), *brinker* (*brk*, BDSC_80438), *asense* (ase, BDSC_44552), *zelda* (*zld*, BDSC_42016), *fruitless* (*fru*, BDSC_66094), *dorsal* (*dl*, BDSC_32934), *Suppressor of Hairless* (*Su(H)*, BDSC_28900) and *daughterless* (*da*, BDSC_38382). Results for *Su(H)* and *da* are shown in Fig. 5F, G and those for the remaining eight factors in Fig. S6A.

### Immunohistochemistry

Larval central nervous systems were dissected in PBS, fixed in 4% paraformaldehyde for 15 min, then antibody staining was carried out as described (Zacharioudaki et al., 2016). The following primary antibodies were used: rat anti-Elav (DSHB; 1:30), rabbit anti-Miranda (a gift from C. Maurange; 1:1000) and goat anti-GFP (Abcam; 1:500). Bound antibodies were visualised with Cy3- and Cy5-conjugated secondary antibodies (Jackson ImmunoResearch; 1:250; Miranda in Cy3, Elav in Cy5), and mNeon was imaged in the green channel. Samples were mounted in VECTASHIELD PLUS Antifade Mounting Medium with DAPI (Vector Laboratories, H-2000-10), which provided the nuclear counterstain.

### Confocal microscopy

Confocal z-stacks were acquired on a Leica SP8 microscope using 40× and 20× oil-immersion objectives (40× for quantification, ≈0.3 µm/pixel; 20× for whole-brain overviews). Four channels were collected: DAPI, GFP reporter, Cy3 (Miranda) and Cy5 (Elav). Images are shown as maximum-intensity projections; scale bars, 50 µm.

### Image analysis and quantification

Reporter fluorescence was quantified from the 40× series using a custom Fiji/ImageJ and Python (readlif, scikit-image) pipeline. For each brain, a central-brain region of interest (ROI) excluding the neuroepithelium/optic lobe, a neuroepithelial (NE) ROI, and a cell-free background ROI were drawn by hand. Neuroblasts (NBs) were detected from the combined GFP and Miranda signal and defined as Miranda-positive cells; GFP was then integrated in three dimensions over each cell with per-slice local background subtraction. Because absolute GFP intensity is sensitive to acquisition settings, absolute NB intensities were compared only within a matched imaging session, and the neuroblast-to-neuroepithelium (NB/NE) GFP ratio was used as a gain-independent, internally normalised readout between sessions. Sample sizes (numbers of cells and of brains per genotype) are reported in the figure legends.

### Sequence and bioinformatic analyses

All genomic coordinates refer to the D. melanogaster dm6 assembly, and analyses were performed with custom Python scripts unless stated otherwise.

#### Cross-species conservation

The short-promoter proximal region (≈−98 to +50 bp relative to the short-promoter TSS, chr2R) was compared across D. melanogaster, D. simulans and D. pseudoobscura. Orthologous sequences were aligned to the D. melanogaster sequence by global Needleman–Wunsch pairwise alignment (match +1, mismatch −1, gap −2), and a position was called conserved when identical across all three species; the Initiator (TCAGT), DPE, the two proximal E-boxes and the proximal motif mutated in the reporter were annotated on the alignment.

#### Core-promoter motif scoring

Candidate sites in the proximal promoter were scored against the JASPAR matrices for Su(H)/CSL (MA0085.2) and the Daughterless/bHLH E-box (MA2308.1). Each position-frequency matrix was converted to a log-odds position-weight matrix (pseudocount 0.5; uniform 0.25 background), and the sense-strand match of the promoter was expressed as a relative score scaled from 0 to 100% between the matrix minimum and maximum; sequence logos display information-content-scaled letter heights. On this scale the proximal motif matched the Daughterless E-box at 86% and a Su(H) site at 74%, and scrambling the element abolished the E-box match.

#### Core-promoter architecture

The interval −100 to +50 bp relative to each annotated TSS was scanned for the canonical core-promoter elements at all five *grh* promoters. An Initiator (Inr; TCAKTY) with a downstream DPE was present in *grh*-short (grh-RN) and *grh*-medium (grh-RK) but absent from the three long promoters, one of which carries GAGA/GAF motifs; no TATA box, BRE, MTE, DRE, Ohler or GC-box element was detected at any of the five (Juven-Gershon and Kadonaga, 2010; Ohler et al., 2002).

#### Genome-wide Initiator (Inr) analysis

Neuroblast- and neuron-“identity” gene sets were derived from neuroblast-versus-neuron RNA-seq (GEO GSE71104) by ranking genes on their neuroblast-versus-neuron log2 fold-change and intersecting the up-regulated sets with sequence-specific DNA-binding transcription factors (Ensembl BioMart, GO:0003700), yielding 107 neuroblast-identity and 134 neuron-identity transcription-factor promoters; a housekeeping set (n = 588) comprised genes expressed stably across neuroblasts, neurons and glia. Promoter sequences were taken around the Ensembl canonical TSS; the Inr ((C/T)A(G/T), or strictly TCAGT) was scored at TSS±2 bp and its position profiled from −15 to +15 bp. Enrichment relative to housekeeping promoters was assessed by Fisher’s exact test, and robustness was confirmed by repeating the analysis on the top 100, 250 and 1000 genes of each set. Because genes that use alternative neuroblast promoters — including the Inr-bearing grh short promoter — are annotated at their long/canonical TSS, they are scored as Inr-negative in this genome-wide analysis, which therefore underestimates Inr usage at alternative stem-cell promoters.

### Published genomic datasets and track visualisation

Genomic tracks were re-analysed from published datasets (dm6/BDGP6): neuroblast ATAC-seq (GEO GSM4561511) and eye-disc ATAC-seq (GEO GSM8186965, series GSE263159); *Su(H)* CUT&RUN in brat-null neuroblast-enriched larval brains (Rajan et al. 2023) (GSE218253); Daughterless ChIP (GEO GSM8561631, from GSE279107; adult head); and neuroblast-versus-neuron RNA-seq (Berger et al. 2012) (GSE38764), shown from the FlyBase L3 CNS neuroblast/neuron tracks. GAF, H3K27ac, H3K14ac and H3K4me1 tracks are from (Zhang et al. 2025) (GSE270072), and the H3K4me3 track is from (Rajan et al. 2023) (GSE218253). For display, signal was read from the published bigWig files over the grh locus (chr2R:17,801,000–17,843,000, dm6) using libBigWig (via the bbi Python bindings) and scaled to each track’s maximum across the locus.

### Quantification and statistical analysis

Graphs were generated in GraphPad Prism 10. Data are plotted as boxes showing the mean ± s.d. with the individual data points overlaid, except in Fig. S6A, where violin plots show the distribution of per-brain ratios with the median and quartiles marked; two-group comparisons used an unpaired two-tailed Student’s t-test (significance thresholds: *P < 0.05; **P < 0.01; ***P < 0.001; ****P < 0.0001). For reporter-intensity graphs each data point is an individual neuroblast, pooled across the brains of a genotype; for the neuroblast-to-neuroepithelium (NB/NE) ratio each data point is one brain, since the ratio is computed per brain. The number of cells and of brains contributing to each comparison is given in the figure legends.

## Acknowledgements

We thank members of the lab for discussions and reagents, M. M. Harrison for the sfGFP-*grh*-long knock-in line, C. Maurange for the anti-Miranda antibody. We used FlyBase (release FB2026_02; Öztürk-Çolak et al., 2024) for gene models, transcript annotations and transcription start sites, for the neuroblast and neuron RNA-seq coverage shown in Fig. 1A, and for information on stocks and alleles. Stocks obtained from the Bloomington Drosophila Stock Center (NIH P40OD018537) were used in this study, including transgenic RNAi lines from the TRiP at Harvard Medical School (NIH/NIGMS R01-GM084947).

## Author contributions

Conceptualisation: S.K. and S.J.B.; Investigation: S.K. and K.M.; Formal analysis: S.K.; Writing – original draft: S.K. S.J.B; Writing – review and editing: S.K., K.M. and S.J.B.; Supervision: S.J.B.

## Data availability

All relevant data are summarised within the paper and Supporting Information. Raw data from quantifications are uploaded to https://doi.org/10.6084/m9.figshare.34329852.

