## Supplementary material for "The Core Promoter has an active role in interpreting patterning information": Suplementary Tables and Figures

### Supplementary Information

**Table S1. Drosophila stocks used in this study.**

| **Stock / line** | **Genotype or description** | **Source (ID)** | **Use** |
|---|---|---|---|
| ***Transgenesis and husbandry*** | | | |
| φC31 / attP40 | *$y^1$ $v^1$ P{nos-phiC31\int.NLS}X; P{CaryP}Msp300$^{attP40}$* | BDSC_25709 | Reporter transgenesis host |
| *nos*-Cas9 | *y1 P(nos-cas9, w+) M(3xP3-RFP.attP)ZH-2A w** | BDSC_54591 | CRISPR injection strain |
| **Endogenous *grh* alleles and knock-ins** | | | |
| *grh*-short::sfGFP | sfGFP knock-in at short (RN/RO) first exon | This study | Short-isoform fusion |
| *grh*-long::sfGFP | sfGFP in first coding exon of long isoforms | Nevil et al. (2020) | Long-isoform fusion |
| *grh*$^{\Delta 10}$ | RN exon-1 deletion; 41 bp junction insertion | This study | Short-promoter LOF |
| *grh*$^{\Delta 15}$ | RN exon-1 deletion; 15 bp junction insertion | This study | Short-promoter LOF |
| *grh*$^{B37}$ | *grh* loss-of-function allele (S.J.B., previously published) | Bray and Kafatos (1991) | Reference allele; Fig S1 cross |
| *Df(2R)Pcl7B* | deficiency uncovering *grh* | Duncan (1982) | Trans-heterozygous test |
| **GAL4 driver and UAS/RNAi lines** | | | |
| *grh*-GAL4 | neuroblast GAL4 driver | Bray lab stock | Drives in NBs |
| UAS-*lacZ* | — | BDSC_4409 | RNAi control |
| UAS-*Taf6*$^{RNAi}$ | TRiP | BDSC_7181 | *Taf6 KD* |

| Stock / line | Genotype or description | Source (ID) | Use |
|---|---|---|---|
| UAS-*Su(H)*$^{RNAi}$ | TRiP | BDSC_28900 | Su(H) KD |
| UAS-*da*$^{RNAi}$ | TRiP.HMS01851 | BDSC_38382 | *da KD* |
| UAS-*klu*$^{RNAi}$ | TRiP | BDSC_28731 | Candidate screen |
| UAS-*br*$^{RNAi}$ | TRiP | BDSC_51189 | Candidate screen |
| UAS-*wor*$^{RNAi}$ | TRiP | BDSC_77364 | Candidate screen |
| UAS-*brk*$^{RNAi}$ | TRiP | BDSC_80438 | Candidate screen |
| UAS-*ase*$^{RNAi}$ | TRiP | BDSC_44552 | Candidate screen |
| UAS-*zld*$^{RNAi}$ | TRiP | BDSC_42016 | Candidate screen |
| UAS-*fru*$^{RNAi}$ | TRiP | BDSC_66094 | Candidate screen |
| UAS-*dl*$^{RNAi}$ | TRiP | BDSC_32934 | Candidate screen |
| **Enhancer–promoter reporter transgenics (this study; *attP40*)** | | | |
| No enhancer | promoter only: pRN | This study | Baseline control |
| *grh[NB]* | × pRN, pRK, pRL, pRJ, pRP, phsp70, pAntp-P2, pRN[mut], pRL[mut] | This study | |
| *dpn[NRE]* | × pRN, pRK, pRL, phsp70 | This study | |
| *βTub56D* | × pRN, pRK, pRL, pRN[mut] | This study | |
| *Act5C* | × pRN, pRK, phsp70, pRN[mut] | This study | |

**Table S2. Promoter and enhancer fragments used in the reporter constructs.**

| Element | Type | Coordinates (dm6) |
|---|---|---|
| *grh*-short (pRN) | promoter, 150 bp (P5; TSS chr2R:17,823,565) | chr2R:17,823,465–17,823,614 |
| *grh*-medium (pRK) | promoter, 150 bp (P4; TSS chr2R:17,814,344) | chr2R:17,814,244–17,814,393 |
| *grh*-long (pRL) | promoter, 150 bp (P3; TSS chr2R:17,802,867) | chr2R:17,802,767–17,802,916 |
| *grh*-long (pRJ) | promoter, 150 bp (P2; TSS chr2R:17,802,246) | chr2R:17,802,146–17,802,295 |
| *grh*-long (pRP) | promoter, 150 bp (P1; TSS chr2R:17,801,132) | chr2R:17,801,032–17,801,181 |
| *hsp70* (phsp70) | minimal promoter, 127 bp (hs43) | chr3R:12,505,750–12,505,876 (Hsp70Bb, plus strand; −43 to +84 of the TSS at chr3R:12,505,793) |
| *Antp-P2* (pAntp-P2) | minimal promoter, 150 bp (Antp P2; TSS chr3R:6,932,840) | chr3R:6,932,791–6,932,940 (minus strand; −100 to +50) |
| *grh*-short[mut] (pRN[mut]) | modified promoter, 150 bp (12 bp replacement) | within chr2R:17,823,465–17,823,614; −88 to −77 (chr2R:17,823,477–17,823,488) |
| *grh*-long[mut] (pRL[mut]) | modified promoter, 150 bp (12 bp replacement) | within chr2R:17,802,767–17,802,916; −88 to −77 |

| | | |
|---|---|---|
| | | (chr2R:17,802,779–17,802,790) |
| *grh[NB]* | enhancer, ≈1.37 kb | chr2R:17,814,846–17,816,211 |
| *dpn[NRE]* | enhancer, ≈411 bp | chr2R:8,236,619–8,237,029 |
| *βTub56D* | enhancer, AS1 region ≈1.36 kb | chr2R:19,449,982–19,451,337 |
| *Act5C* | enhancer, ≈1.66 kb (*HindIII*–*SpeI* fragment) | chrX:5,898,687–5,900,342 |

**Table S3. Full genotypes by figure panel.**

| **Figure panel** | **Genotype** |
|---|---|
| Fig. 1C, S1A | *grh*-RL::sfGFP knock-in / + ; *grh*-RN::sfGFP knock-in / + (imaged separately) |
| Fig. S1B | $grh^{\Delta15}$ / $grh^{\Delta15}$ ; $grh^{\Delta10}$ / $grh^{\Delta10}$ ; wild type (w[*]) |
| Fig. S1C | $grh^{\Delta15}$ or $grh^{\Delta10}$ over $grh^{B37}$, and *Df(2R)Pcl7B,* against CyO |
| Fig. 2B,C | w[*] ; P{*grh-NB*-pRN}attP40 / + (equivalently *grh-NB*-pRK, *grh-NB*-pRL, *grh-NB*-pRJ, *grh-NB*-pRP) |
| Fig. 2D | w[*] ; P{*grh-NB*-phsp70}attP40 / + and w[*] ; P{*grh-NB*-pAntp-P2}attP40 / + |
| Fig. 3B,C,E | w[*] ; P{*dpn-NRE*-pRN}attP40 / + (equivalently pRK, pRL) |
| Fig. 3D | w[*] ; P{*dpn-NRE*-phsp70}attP40 / + |

| | |
|---|---|
| Fig. 4B | w[*] ; P{*βTub56D*-pRN}attP40 / + (equivalently *βTub56D*-pRK, *βTub56D*-pRL) |
| Fig. 4D | w[*] ; P{*Act5C*-pRN}attP40 / + (equivalently *Act5C*-pRK, *Act5C*-phsp70) |
| Fig. 5B,C | w[*] ; P{*βTub56D*-pRN}attP40 / + ; *grh*-GAL4 / UAS-*Taf6*[RNAi] (BDSC 57181) — control UAS-lacZ |
| Fig. 5D,E | w[*] ; P{*grh-NB*-pRN}attP40 / + and w[*] ; P{*grh-NB*-pRN[mut]}attP40 / + |
| Fig. 5F,G | w[*] ; P{*βTub56D*-pRN}attP40 / + ; *grh*-GAL4 / UAS-*da*[RNAi] (BDSC 38382, TRiP.HMS01851) and / UAS-*Su(H)[*RNAi] (BDSC 1208) — control UAS-lacZ |
| Fig. S2 | w[*] ; P{*grh-NB*-pRN}attP40 / + versus w[*] ; P{pRN}attP40 / + (promoter only, no enhancer) |
| Fig. S3 | w[*] ; P{*grh-NB*-pRN}attP40 / + ; P{*dpn-NRE*-pRN}attP40 / + ; P{*Act5C*-pRN}attP40 / + , each also with phsp70 |
| Fig. S4 | w[*] ; P{*βTub56D*-pRN}attP40 / + , *βTub56D*-pRK, *βTub56D*-phsp70 ; P{*Act5C*-pRN}attP40 / + , *Act5C*-pRK, Act5C-phsp70 |
| Fig. S5A–D | w[*] ; P{*βTub56D*-pRN}attP40 / + versus P{*βTub56D*-pRN[mut]}attP40 / + ; and *Act5C*-pRN versus *Act5C*-pRN[mut] |
| Fig. S5E,F | w[*] ; P{*grh-NB*-pRL}attP40 / + versus w[*] ; P{*grh-NB*-pRL[mut]}attP40 / + |
| Fig. S6A | w[*] ; P{*βTub56D*-pRN}attP40 / + ; *grh*-GAL4 / UAS-X[RNAi], X = *klu* (BDSC 28731), *br* (51189), *wor* (77364), *brk* (80438), *ase* (44552), *zld* (42016), *fru* (66094)— control UAS-lacZ |

| | |
|---|---|
| Fig. S6B,C | *ms1096*-GAL4 / + ; + ; Su(H)-GFP / UAS-*Su(H)[*RNAi] (BDSC_25900) — control, Su(H)-GFP / UAS-lacZ. |

### Supplementary figures and legends

**Figure S1. Complementary isoform expression in imaginal discs and the short-isoform requirement for neuroblast elimination.** (A) Endogenous GFP for the long- and short-isoform knock-ins in different imaginal discs. (B) Larval CNS of wild type, *grhΔ15*/*grhΔ15* and *grhΔ10*/*grhΔ10* stained for Miranda (Mira); both frameshift mutants retain supernumerary abdominal ventral-nerve-cord neuroblasts (boxed). (C) Viability of the indicated crosses (*grhΔ15* and *grhΔ10* over *grhB37*, and *Df(2R)Pcl7B*), scored by *CyO* balancer and sex (males, females). Maximum-intensity projections; scale bars, 50 µm. Full genotypes are given in Table S3.

**Figure S2. The short promoter has no autonomous, enhancer-independent activity.** (A) Expression from *grh[NB]* in combination with *grh*-short promoter (grh[NB]≫short RN prom) in comparison to *grh*-short (RN) promoter alone. The promoter alone yields no detectable expression (GFP, Maximum-intensity projections, inverted greyscale; scale bars, 50 µm.). (B) Quantification (background-subtracted GFP relative to local background) of neuroblast reporter intensity, grh[NB]≫pRN versus pRN alone. (n = 4 brains per genotype). Scatter plot showing the mean ± s.d.,. Full genotypes are given in Table S3.

**Figure S3. Promoter competence is cell-type-dependent: enhancer×promoter combinations in the eye–antennal disc.** Third-instar eye–antennal discs carrying *grh*[NB], *dpn[NRE]* or *Act5C* enhancer paired with *grh*-short (RN) or *hsp70* promoters(GFP, inverted greyscale). The *grh*[NB] is inactive in this tissue with either promoter; *dpn[NRE]* drives expression in the photoreceptor field with *grh*-short (RN) or *hsp70* promoters comparably; in contrast to the larval CNS (Figure 4), *Act5C* enhancer drives strong expression with *hsp70* promoter not with *grh*-short promoter . Maximum-intensity projections, inverted greyscale; scale bars, 50 µm. Full genotypes are given in Table S3.

**Figure S4. Promoter selectivity by ubiquitous enhancers in the wing disc.** Expression in third-instar wing imaginal discs from *βTub56D* enhancer (top) or *Act5C* enhancer (bottom) combined with *grh*-short (RN), *grh*-med (RK) or *hsp70* promoters. Maximum-intensity projections; scale bars, 50 µm. Full genotypes are given in Table S3.

**Figure S5. Effects from mutating the proximal motif .** (A,B) Expression from the *βTub56D* enhancer paired with wild type *grh*-short (RN) versus motif-mutated (RN[mut]) (GFP, inverted greyscale), with quantification of neuroblast reporter intensity (B). (n=3 brains for each genotype) (C,D) Expression from *Act5C* enhancer paired with wild-type *grh*-short versus motif-mutated promoter, with quantification of neuroblast reporter intensity (D). (n = 3 brains for each genotype) (E,F) Expression from *grh[*NB] enhancer combined with wild-type *grh*-long (RL) promoter versus modified *grh*-long with 12 bp motif at the equivalent position (n=3 brains for each genotype). Maximum-intensity projections; scale bars, 50 µm. Data are plotted as boxes showing the mean ± s.d., with individual neuroblasts overlaid; groups were compared with an unpaired two-tailed Student's t-test (n.s., not significant; *P < 0.05; **P < 0.01; ***P < 0.001; ****P < 0.0001). Full genotypes are given in Table S3.

**Figure S6. A candidate screen for factors acting on the *grh*-short promoter**

(A) The proximal element scored against JASPAR matrices: it matches the Su(H)/CSL motif (MA0085.2) at ≈74% and the Daughterless/bHLH E-box (MA2308.1) at ≈86%.

(B) Neuroblast-to-neuroepithelium (NB/NE) GFP ratio for the βTub56D≫*grh*-short reporter after neuroblast-restricted RNAi (*grh*-GAL4) against the indicated candidates (*klu,* *br, wor, brk, ase, zld,* and *fru*), each compared with the *UAS-lacZ* control. Violin plots show the distribution of per-brain NB/NE ratios with the median and quartiles marked, and individual brains overlaid (*UAS-lacZ* control, n = 6 brains; n = >4 brains per RNAi genotype). (C,D) Validation of the *Su(H)* RNAi: *ms1096*-GAL4 driving *UAS-Su(H)^RNAi* in the context of Su(H)::GFP transgene. Su(H)::GFP levels are depleted in the wing pouch relative to control, confirming effective knockdown. In (D) the ratio of Su(H)::GFP in the pouch to background is plotted as a box showing the mean ± s.d.; control n = 7 wing discs and *Su(H)* RNAi n = 6. Scale bars, 50 µm. Groups were compared with an unpaired two-tailed Student's t-test (P value < 0.0001). Full genotypes are given in Table S3.

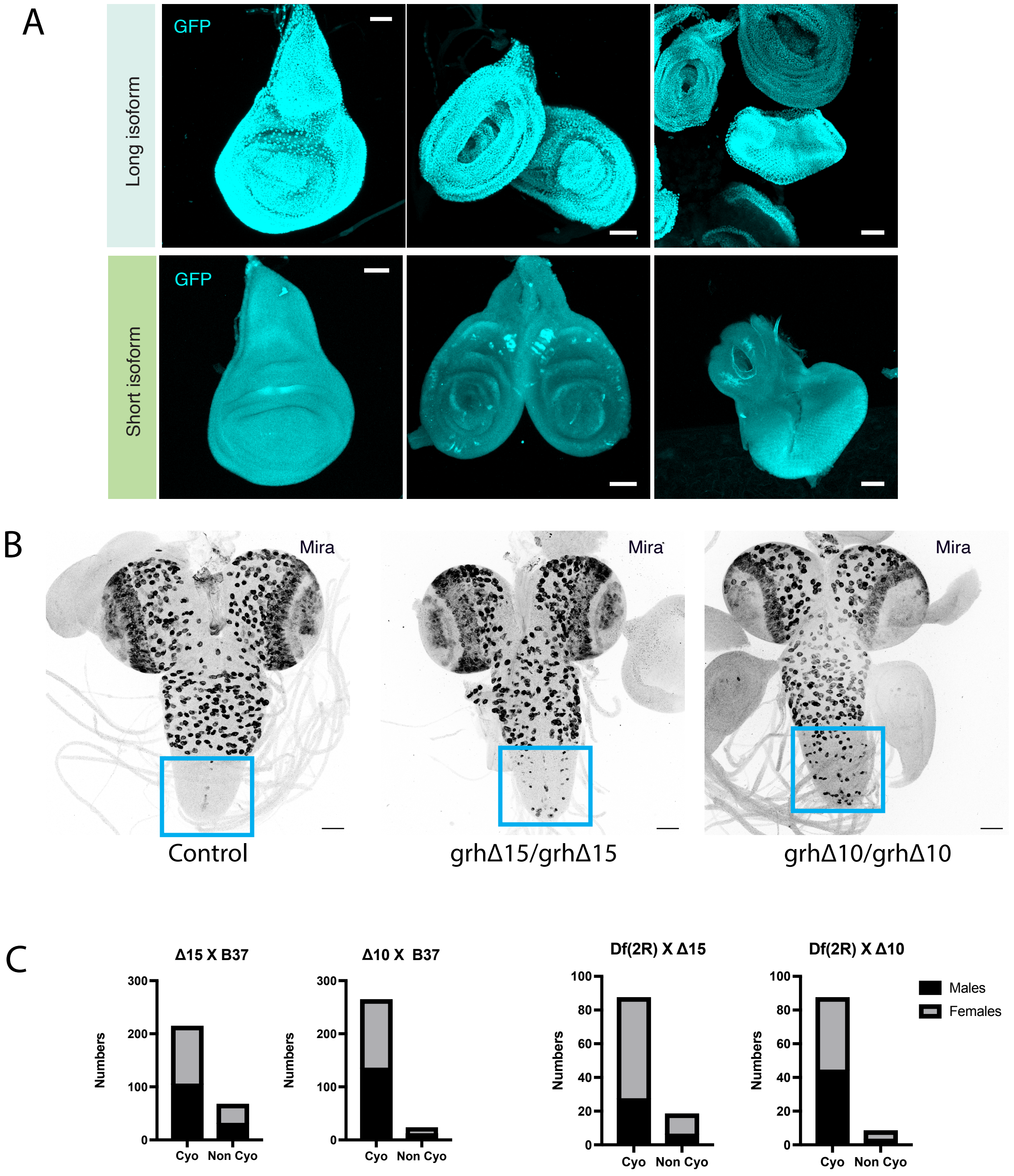


**Figure S1. Complementary isoform expression in imaginal discs and the short-isoform requirement for neuroblast elimination.** (A) Endogenous GFP for the long- and short-isoform knock-ins in different imaginal discs. (B) Larval CNS of wild type, *grhΔ15/grhΔ15* and *grhΔ10/grhΔ10* stained for Miranda (Mira); both frameshift mutants retain supernumerary abdominal ventral-nerve-cord neuroblasts (boxed). (C) Viability of the indicated crosses (*grhΔ15* and *grhΔ10* over *grhB37*, and *Df(2R)Pcl7B*), scored by *CyO* balancer and sex (males, females). Maximum-intensity projections; scale bars, 50 µm. Full genotypes are given in Table S3.

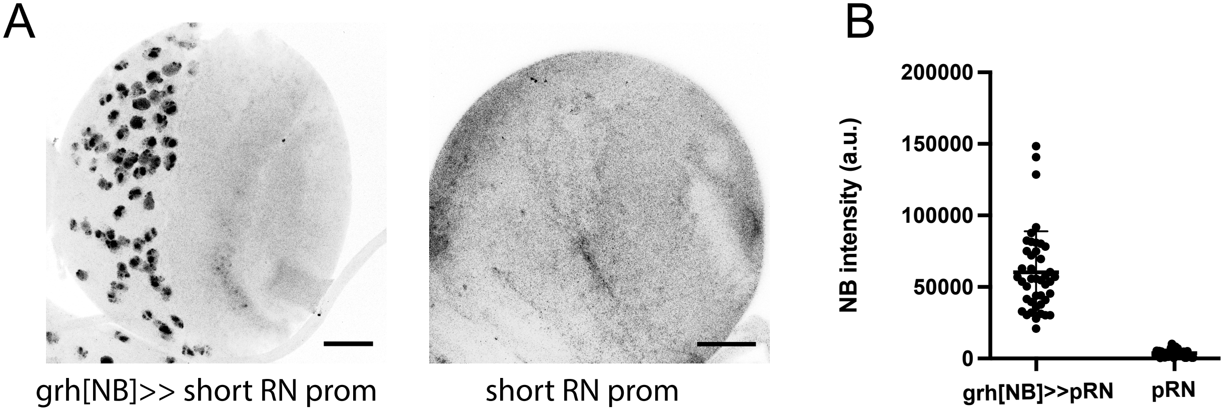


**Figure S2. The short promoter has no autonomous, enhancer-independent activity.** (A) Expression from *grh[NB]* in combination with *grh*-short promoter (grh[NB]▯short RN prom) in comparison to *grh*-short (RN) promoter alone. The promoter alone yields no detectable expression (GFP, Maximum-intensity projections, inverted greyscale; scale bars, 50 µm.). (B) Quantification (background-subtracted GFP relative to local background) of neuroblast reporter intensity, grh[NB]▯pRN versus pRN alone. (n = 4 brains per genotype). Scatter plot showing the mean ± s.d.,. Full genotypes are given in Table S3.

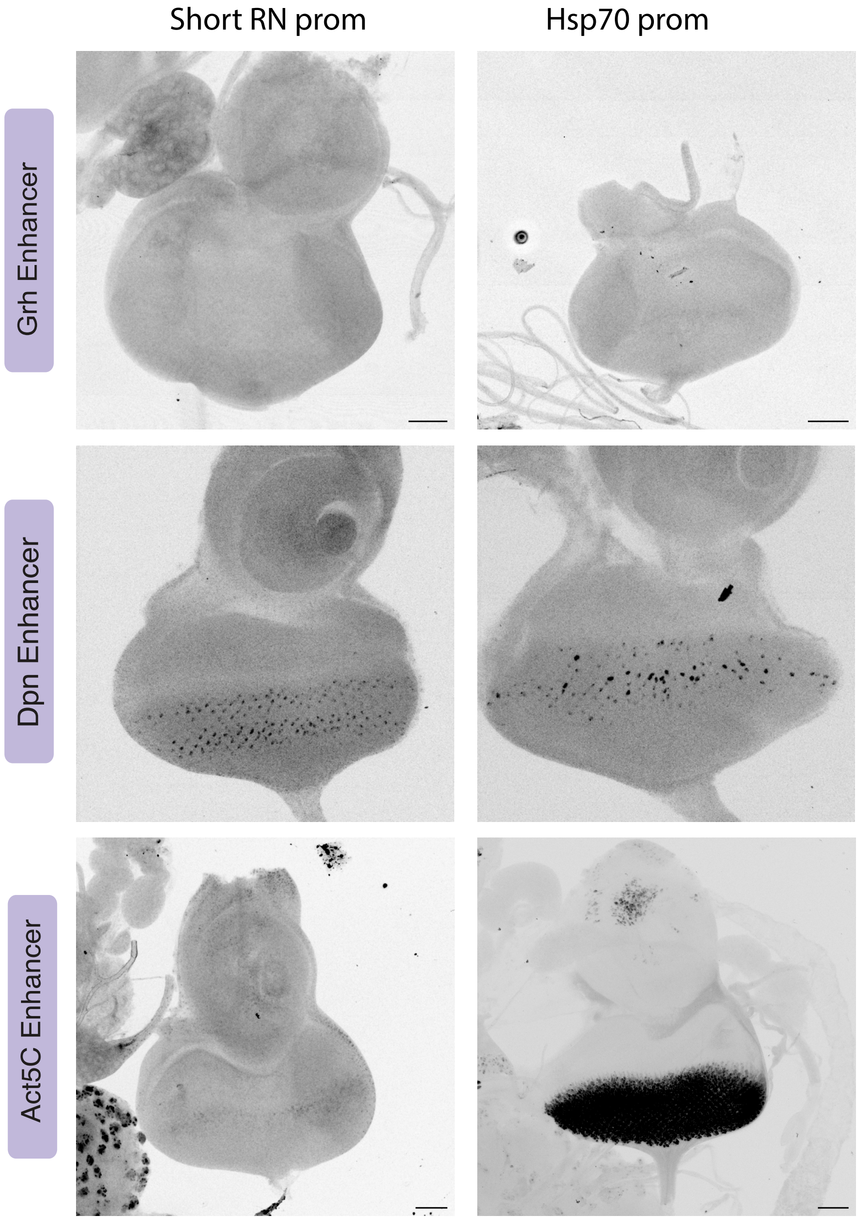


**Figure S3. Promoter competence is cell-type-dependent: enhancer×promoter combinations in the eye–antennal disc.** Third-instar eye–antennal discs carrying *grh*[NB], *dpn[NRE]* or *Act5C* enhancer paired with *grh*-short (RN) or *hsp70* promoters(GFP, inverted greyscale). The *grh*[NB] is inactive in this tissue with either promoter; *dpn[NRE]* drives expression in the photoreceptor field with *grh*-short (RN) or *hsp70* promoters comparably; in contrast to the larval CNS (Figure 4), *Act5C* enhancer drives strong expression with *hsp70* promoter not with *grh*-short promoter . Maximum-intensity projections, inverted greyscale; scale bars, 50 μm. Full genotypes are given in Table S3.

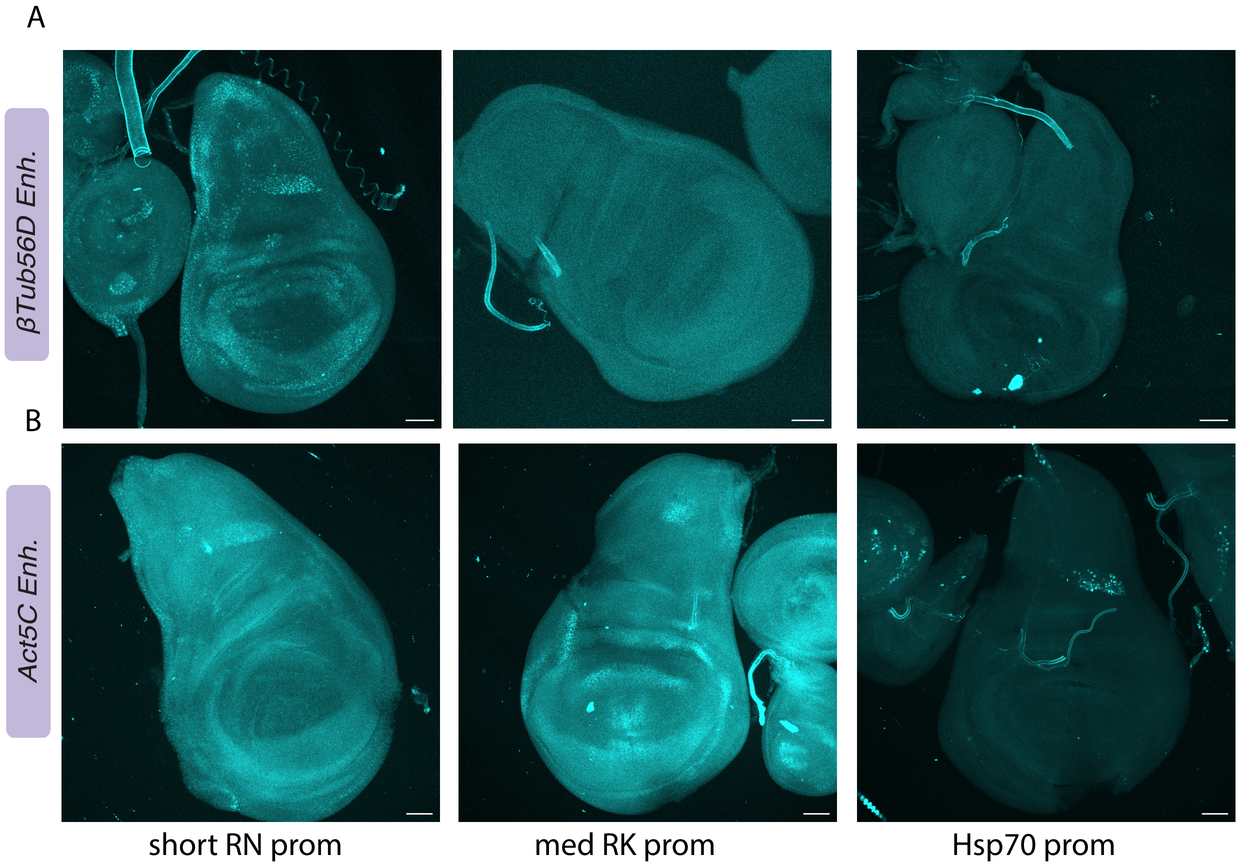


**Figure S4. Promoter selectivity by ubiquitous enhancers in the wing disc.** Expression in third-instar wing imaginal discs from *βTub56D* enhancer (top) or *Act5C* enhancer (bottom) combined with *grh*-short (RN), *grh*-med (RK) or *hsp70* promoters. Maximum-intensity projections; scale bars, 50 µm. Full genotypes are given in Table S3.

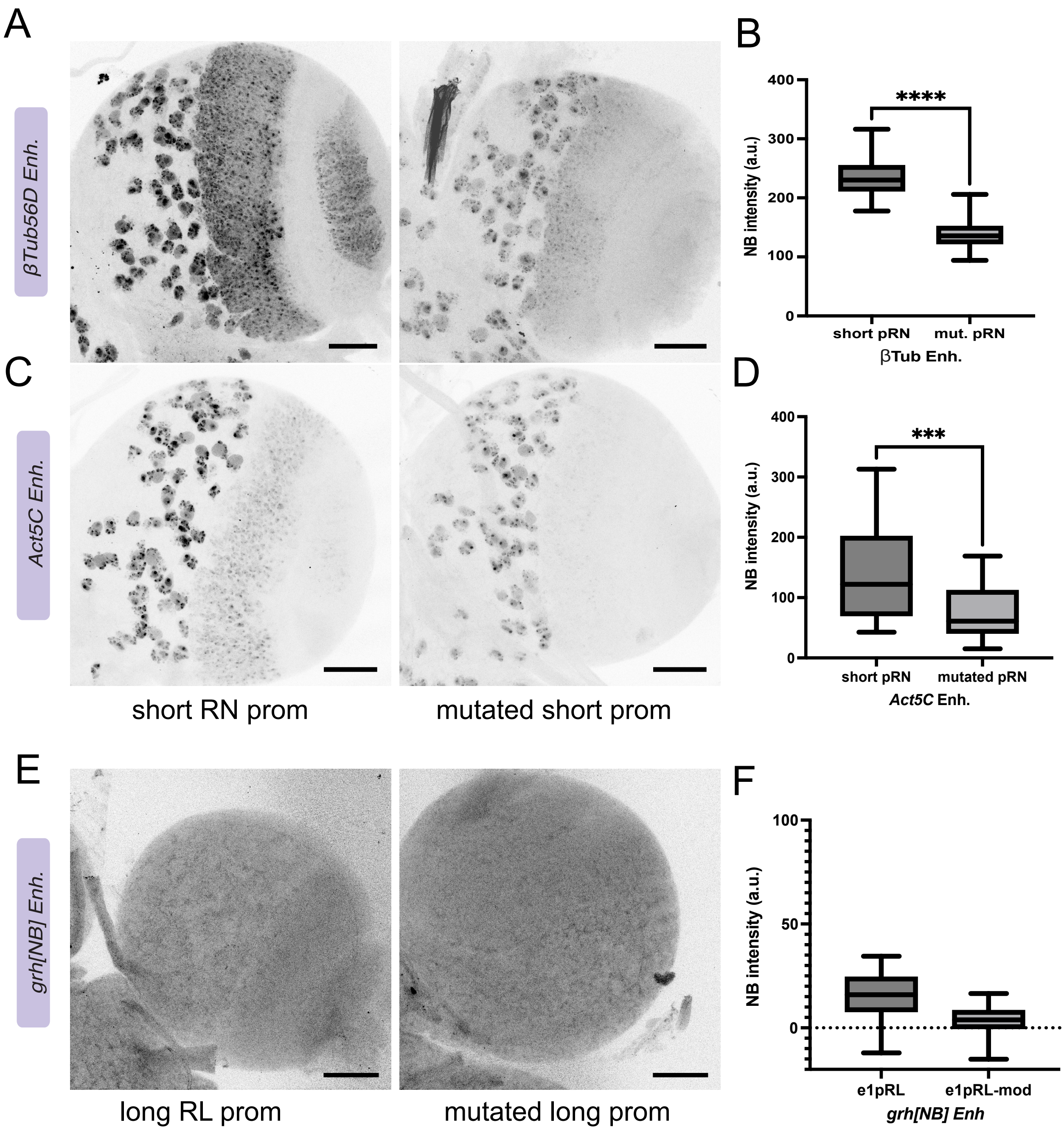


**Figure S5. Effects from mutating the proximal motif .** (A,B) Expression from the *βTub56D* enhancer paired with wild type *grh*-short (RN) versus motif-mutated (RN[mut]) (GFP, inverted greyscale), with quantification of neuroblast reporter intensity (B). (n=3 brains for each genotype) (C,D) Expression from *Act5C* enhancer paired with wild-type *grh*-short versus motif-mutated promoter, with quantification of neuroblast reporter intensity (D). (n = 3 brains for each genotype) (E,F) Expression from *grh[*NB] enhancer combined with wild-type *grh*-long (RL) promoter versus modified *grh*-long with 12 bp motif at the equivalent position (n=3 brains for each genotype). Maximum-intensity projections; scale bars, 50 µm. Data are plotted as boxes showing the mean ± s.d., with individual neuroblasts overlaid; groups were compared with an unpaired two-tailed Student's t-test (n.s., not significant; *P < 0.05; **P < 0.01; ***P < 0.001; ****P < 0.0001). Full genotypes are given in Table S3.

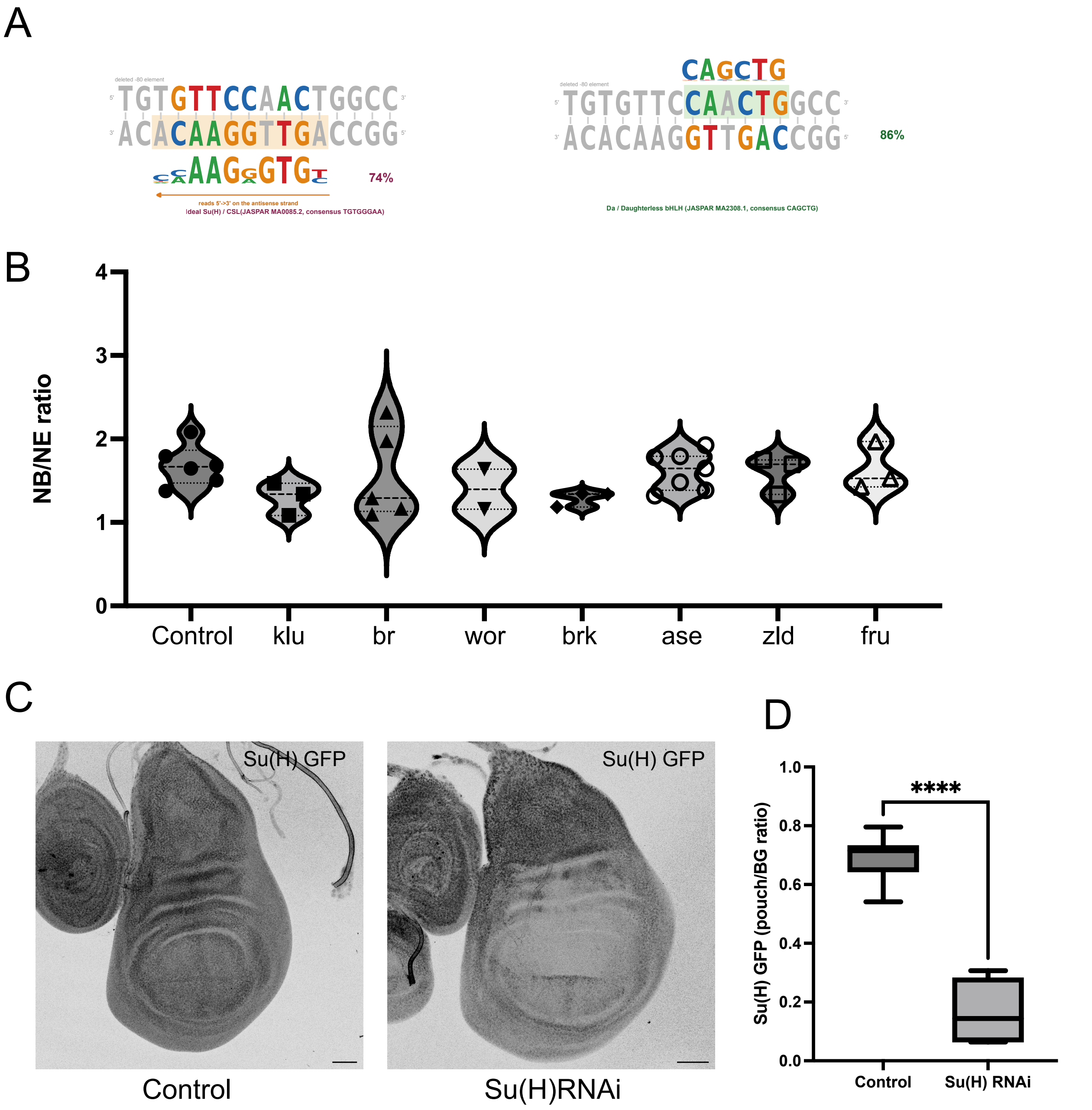


**Figure S6. A candidate screen for factors acting on the *grh*-short promoter** (A) The proximal element scored against JASPAR matrices: it matches the Su(H)/CSL motif (MA0085.2) at ≈74% and the Daughterless/bHLH E-box (MA2308.1) at ≈86%. (B) Neuroblast-to-neuroepithelium (NB/NE) GFP ratio for the βTub56D⊠*grh*-short reporter after neuroblast-restricted RNAi (*grh*-GAL4) against the indicated candidates (*klu, br, wor, brk, ase, zld,* and *fru*), each compared with the *UAS-lacZ* control. Violin plots show the distribution of per-brain NB/NE ratios with the median and quartiles marked, and individual brains overlaid (*UAS-lacZ* control, n = 6 brains; n = >4 brains per RNAi genotype). (C,D) Validation of the *Su(H)* RNAi: *ms1096*-GAL4 driving *UAS-Su(H)^RNAi* in the context of Su(H)::GFP transgene. Su(H)::GFP levels are depleted in the wing pouch relative to control, confirming effective knockdown. In (D) The ratio of Su(H)::GFP in the pouch to background is plotted as a box showing the mean ± s.d.; control n = 7 wing discs and *Su(H)* RNAi n = 6. Scale bars, 50 µm. Groups were compared with an unpaired two-tailed Student's t-test (P value < 0.0001). Full genotypes are given in Table S3.
